# Niche differential gene expression analysis in spatial transcriptomics data identifies context-dependent cell-cell interactions

**DOI:** 10.1101/2023.01.03.522646

**Authors:** Kaishu Mason, Anuja Sathe, Paul Hess, Jiazhen Rong, Chi-Yun Wu, Emma Furth, Hanlee P. Ji, Nancy Zhang

**Affiliations:** Department of Statistics and Data Science, The Wharton School, University of Pennsylvania; Division of Oncology, Department of Medicine, Stanford University School of Medicine, Stanford, CA, United States; Genomics and Computational Biology Graduate Program, Perelman School of Medicine, University of Pennsylvania; The Gladstone Institute; Department of Pathology and Laboratory Medicine, Perelman School of Medicine, University of Pennsylvania

## Abstract

Single cells influence, and are shaped by, their local spatial niche. Technologies for in situ measurement of gene expression at the transcriptome scale have enabled the detailed profiling of the spatial distributions of cell types in tissue as well as the interrogation of local signaling patterns between cell types [1]. Towards these goals, we propose a new statistical procedure called niche-differential expression (niche-DE) analysis. Niche-DE identifies cell-type specific niche-associated genes, defined as genes whose expression within a specific cell type is significantly up- or down-regulated, in the context of specific spatial niches. We develop effective and interpretable measures for global false discovery control and show, through the analysis of data sets generated by myriad protocols, that the method is robust to technical issues such as over-dispersion and spot swapping. Niche-DE can be applied to low-resolution spot- and ROI-based spatial transcriptomics data as well as data that is single-cell or subcellular in resolution. Based on niche-DE, we also develop a procedure to reveal the ligand-receptor signaling mechanisms that underlie niche-differential gene expression patterns. When applied to 10x Visium data from liver metastases of colorectal cancer, niche-DE identifies marker genes for cancer-associated fibroblasts and macrophages and elucidates ligand-receptor crosstalk patterns between tumor cells, macrophages and fibroblasts. Co-detection by indexing (CODEX) was performed on the same patient samples, to corroborate the niche-DE results.

## Introduction

Spatial transcriptomic (ST) technologies such as Merfish [2], SeqFISH+ [3], Visium [4], Slide-seq [5], and Stereo-seq [6] are rapidly maturing, enabling the high resolution in situ measurement of gene expression at the transcriptome scale. To mine this data, computational methods have been proposed for image segmentation [7], technical artifact removal [8], spot deconvolution for spot-based data [9], spatially variable gene detection [10], and other downstream analyses [1]. In particular, the detection of spatially variable genes, defined as genes displaying clear spatial patterns in expression, has become a standard analysis step. Spatially variable genes can be used to visualize tissue architecture and identify cell types that have distinct spatial localization. Once the distribution of cell types have been determined at a macro-level, it is often of interest to interrogate local, cell-type specific interactions [11]. By design, spatially variable gene analysis is geared towards the identification of global patterns in gene expression, not local interactions between cell types. For example, in cancer tissue, spatially variable genes can easily demarcate cancer vs normal tissue regions. However, it is unclear how to identify local patterns such as type-specific niche signaling between tumor and immune cells.

Local interactions between cells, such as those based on signaling pathways or chemical and biomechanical remodeling of the extracellular matrix, are highly cell type specific. Thus, we explore cell-type-specific models for the dependence of a cell’s gene expression program on its local microenvironment niche. To this end we propose a new statistical procedure called niche-differential expression (niche-DE) analysis. Niche-DE identifies cell-type specific niche-associated genes, defined as genes whose expression within specific cell type(s) is significantly up- or down-regulated in the context of specific spatial niches, as compared to their cell-type-specific mean expression. Although niche-DE is conceptually defined at the single-cell level, we derive an equivalent model for the recovery of niche-DE genes from lower resolution spatial transcriptomic data where each observation is a spot (or region of interest) containing a mixture of cell types. To ensure rigorous and reproducible analyses, we propose an interpretable framework for FDR control that reports significant signals at three levels (gene, cell type, and interaction level). Through simulations, we show that the method is robust to overdispersion and spot swapping.

One attractive feature of niche-differential expression analysis versus spatially variable gene analysis is that the former is naturally amenable to cross-sample integration. A statement such as “Macrophages upregulate expression of a given gene *X* when situated near tumor cells” can be applied and tested across multiple tissue slices. In contrast, a statement such as “*X* is a spatially variable gene that is upregulated in the left corner of the tissue” cannot be directly applied across samples due to differences in frames of reference. As such, existing analyses of spatial transcriptomic data perform spatially variable gene detection on each tissue slice separately, leaving open the question of how these genes could be interpreted across slices.

When niche-differentially expressed genes are identified, a natural follow-up inquiry is whether a known inter-cell signaling mechanism is responsible for their up- or down-regulation. Towards this goal, we integrate niche-DE statistics with Niche-net ligand-target matrices [11] to identify ligand-receptor signaling channels between cell types. We also develop rigorous test statistics for this step, and show that they can recover cell-cell signaling interactions for spot-based spatial transcriptomic data where each spot may be a mixture of cells.

We illustrate niche-DE through the analysis of 10x Visium data consisting of liver metastases of colorectal cancer from 5 patients. Niche-DE identifies known and novel marker genes for tumor-associated fibroblasts and macrophages and elucidates ligand-receptor crosstalk patterns between tumor cells, macrophages and fibroblasts. These findings are corroborated by parallel analyses of single cell RNA sequencing (scRNA-seq) and CODEX data for the same tissue type, and in one case, from the same patient.

## Results

### Single-cell niche-differential gene expression model

We start by describing the niche-DE model, first for data of single-cell resolution and then for data of lower-resolution. We then summarize the framework for FDR control and bandwidth selection. Details of the algorithm are given in the Methods section.

For spatial transcriptomic data at *single cell* resolution, we assume that the cells have already been labeled by type, e.g. through the use of marker genes or a reference-based label transfer method [12, 13]. Let *Y_c,g_* be the observed count for gene *g* ∈ {1,..., *G*} in cell *c* ∈ {1,..., *C*}, *T_c_* ∈ {1,…, *T*] be the type of cell *c*, and *μ_t,g_* = *E*(*Y_c,g_*|*T_c_* = *t*) be the expected expression of gene *g* in a cell of type *t*. For each cell, we summarize its spatial neighborhood through the computation of a kernel-smoothed density of cell type compositions centered at the cell (Fig. 1A):

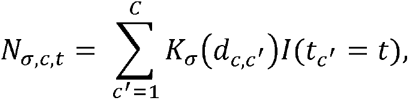

where *d_c,c_*′ is the physical distance between cells *c*, *c*′ and *K_σ_*(*d*) is a kernel of bandwidth *σ*, e.g. 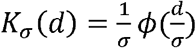 where *ϕ* is the Gaussian filter. We call *N_σ,c_* = {*N*_*σ,c*,1_,…, *N_σ,C,T_*) the effective niche cell type composition of cell *c* with kernel bandwidth *σ*. In defining *N_σ,c_*, we refer to cell *c* as the index cell (Fig. 1A). We consider first the case where the bandwidth *σ* is given, and later discuss the pooling of evidence across multiple *σ*.

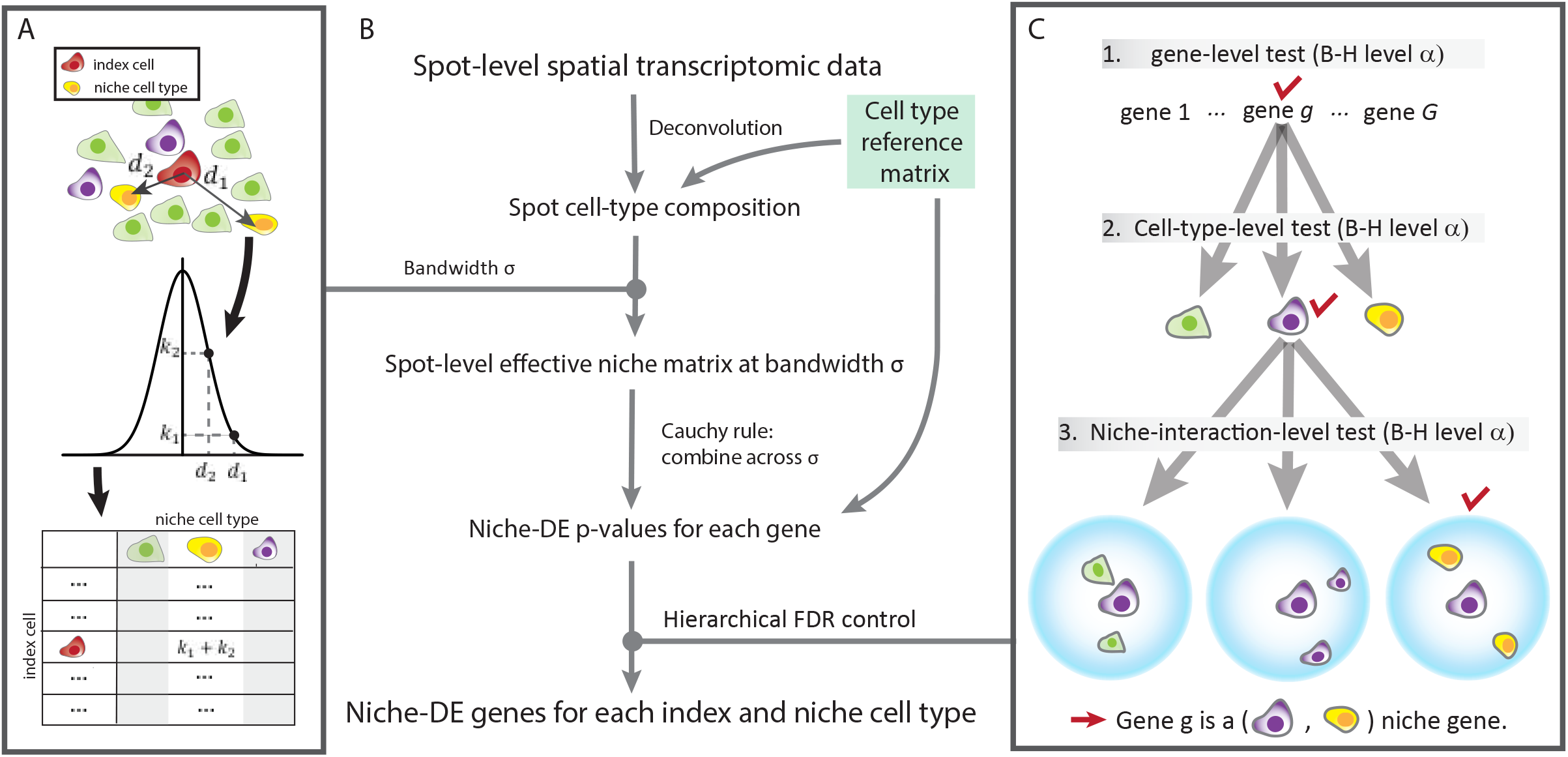
1A) Schematic of effective niche calculation: We aim to quantify the cell type composition of each cell’s neighborhood. For each index cell, we calculate the pairwise kernel distance similarity between itself and each other cell in the sample. We use a Gaussian kernel with bandwidth *σ*. The effective niche for the index cell is a vector with dimension equal to the number of unique cell types in the sample where index *i* represents the sum of kernel similarities between the index cell and the cells of type *i*. 1B) Schematic of niche-DE pipeline: To perform niche-DE, we first perform deconvolution/cell type identification of our data. We then calculate the effective niche using a Gaussian kernel of bandwidth *σ*. We then apply the regression based niche-DE model using the effective niche calculated in the previous step. If one desires, they can repeat this step for multiple bandwidths. Using the Cauchy combination test across the different kernels used, we can calculate a pvalue for testing whether gene *g* is an (*i, n*) niche gene for all genes *g* and index-niche cell type pairs. 1C) FDR control: To guarantee correct FDR control, we utilize the hierarchical Benjamini-Hochberg procedure[14]. We first test if a gene shows evidence of being a niche-DE gene in any index-niche pair. This results in a pvalue for each gene. We then apply the BH procedure to this set of genes. For a gene whose adjusted pvalue is below the cutoff value, we test if it a niche gene in with index cell type *i*. Testing across all *T* unique cell types in our sample, we get *T* celltype specific p-values for each gene *g* that is tested at this level. We then apply the Benjamini-Hochberg correction at level *α* across these pvalues for each gene. For all gene, index-cell types pairs (*g, i*) that are significant after correction, we proceed to test if gene *g* is an (*i,n*) niche gene for each niche cell type *n*. After applying the Benjamini-Hochberg correction at level *α* across all *T* pvalues for each (*g, i*) pair, if a (*g, i, n*) set has a pvalue below the cutoff value, we conclude that gene *g* is an (*i,n*) niche gene.

In the single-cell niche differential gene expression model, the expression of gene *g* in index cell *c* is Negative-Binomial-distributed with dispersion *γ_g_* and mean that varies according to both the index cell type and the effective niche:

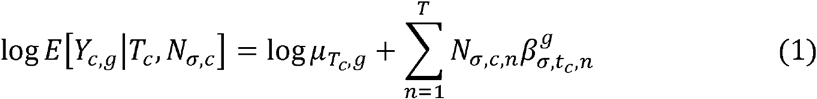

where *μ_T_c_,g_* = *E*[*Y_c,g_*|*T_c_*] is the global mean of gene *g* in cells of type *T_c_*. Let *i* ∈ {1,…, *T*] be the index cell type, and *n* ∈ {1,…, *T*} be the niche cell type, Our goal is to estimate the cell-type specific niche-differential expression parameters 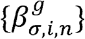, and to identify those genes *g* and index-niche cell type combinations (*i, n*) where 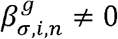. A significant test against the null hypothesis 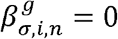 means that, when the index cell is of type *i*, the enrichment of cell type n in the effective niche is associated with a significant change in the expression of gene *g* within the index cell. In this case, we call gene *g* a (*i, n*)^+^ niche gene if the association is positive and a (*i, n*)^-^ niche gene if the association is negative. Note that the relationship between niche and index cell type is asymmetric.

### Detecting niche-differential expression from spot-level data

Now consider the case where, instead of single cells, we observe spots *s* ∈ {1,…, *S*}, where each spot may be a mixture of cells of different types. Let *X_s,g_* be the count of gene *g* in spot *s*. In Methods, we show that model (1) is approximately equivalent to the following spot-level model:

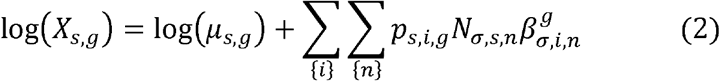

where *μ_s,g_* = ∑_*c:s*(*c*)=*s*_ *μ_T_c_,g_* is the expected expression of gene *g* in spot s given the true cell type composition of spot *s*, *p_s,i,g_* = (*μ_s,g_*)^-1^ ∑_*c:s*(*c*)=*s*_ *μ_t_c_,g_I*(*t_c_* = *i*) is the expected proportion of gene *g*’s expression in spot *s* that originate from cell type *i*, and *N_σ,s,n_* is the concentration of cell type n in the effective niche of spot *s*. As shown in Fig. 1B, we start by computing the terms *μ_s,g_*, *p_s,i,g_* and *N_σ,s,n_* via spot deconvolution [9] Then, the terms 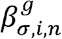 can be estimated along with their corresponding pvalues 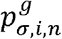 via negative binomial regression. It is important to note that, despite the fact that model (2) is fit using lower-resolution spot-level data, due to the set-up of the model, the interpretation of the parameters 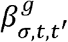 are exactly the same as for model (1).

### Hierarchical False discovery rate control

For each gene, hypothesis tests are performed for *T*^2^ parameters: one parameter 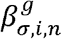 for each index-niche cell type pair. Across *G* genes, this amounts to a parallel screen of *GT*^2^ p-values. As shown in Fig. 1C, we control the false discovery rate (FDR) through a hierarchical Benjamini Hochberg procedure [14]. Specifically, we start with a gene-level test, to identify genes that are niche-DE in *at least one* of the *T*^2^ possible cell-type configurations. This is achieved through applying the Cauchy combination procedure with uniform weights on the set of pvalues obtained through testing the null hypothesis 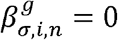 for all *i, n* [15]. A Benjamini-Hochberg correction at level *α* is applied to the gene-level p-values. For genes that are rejected at this level, we proceed to test at the cell type level: for each cell type *i*, we apply the Cauchy combination procedure on the set of pvalues obtained through testing the null hypothesis 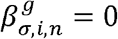 for all niche cell types *n*. Thus, we get *T* cell-type specific p-values for each gene *g* that is tested at this level, and Benjamini-Hochberg correction at level *α* is applied across genes. For all gene and index-cell types pairs (*g, i*) that are significant after correction, we proceed to test whether 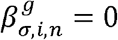 for each niche cell type *n*, again applying a Benjamini-Hochberg correction at level *α*.

This hierarchical procedure can be shown to control overall FDR control at approximate level *α*(#*Discoveries + #Families Tested*)/(#*Discoveries* + 1). Importantly, this procedure allows the flexibility to report niche effects at the gene and index-cell type levels when, due to multicollinearity caused by co-localizing cell types, it may be difficult to tease apart which cell type in the niche is responsible for the niche-differential expression. Such colocalization induced multicollinearity can be rampant even when the data is at single-cell resolution. In the case where a gene rejects at the cell type level for cell type *i* but is not a (*i, n*) niche gene for any niche cell type *n*, we call this gene an *i*-niche gene. Note that it is also possible to screen for one-sided niche effects by testing the null hypothesis 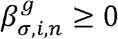 or 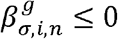.

### Applying niche-DE over multiple kernel bandwidths

The kernel bandwidth parameter *σ* determines the spatial range of cells that contribute towards the effective niche of the index cell/spot. Because we do not know the optimal *σ* a priori, we perform niche-DE on a grid of *K* different kernels *σ*_1_,…. *σ_K_*. This gives us gene-level p-values 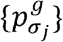 for each bandwidth *σ_j_*. We also record the log-likelihood score, 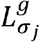, of the negative binomial regression of *X*_{_*s,g*_}_, on *N_σ,s_*. To pool together pvalues from each kernel, we use the Cauchy combination test with the weight for 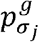. defined as 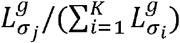 to get a kernel weighted gene level p-value *p^g^*. We apply the BH procedure on the set of kernel-weighted pvalues *p^g^* to determine which genes proceed to the cell type level test. This procedure allows us to test multiple kernel bandwidths, prioritizing bandwidths that give better fit to the data. The same bandwidth-weighting procedure is performed at the cell type and interaction levels.

### Precision of false positive control and robustness to spot swapping

First and foremost, we evaluate the accuracy of type I error control for niche-DE testing to ensure that false positives are controlled in the transcriptome-level screen across cell-type configurations. Towards this end, we generated realistic spatial transcriptomic data in the absence of niche-DE genes, as shown in Fig. 2A: Starting from a high-quality Liver mCRC Visium data set matched with scRNA-seq data, we deconvolved each spot s to estimate its cell type composition vector, which, when multiplied with the reference cell-type specific gene expression matrix, gave us the expected expression vector for that spot *μ_s_*. We then sampled a new expression vector *X_s_* for the spot by drawing from a negative binomial distribution with mean *μ_s_* and a given overdispersion parameter. Note that, since expression values are drawn for each spot independently given only the spot’s cell type decomposition and without influence from its neighbors, data simulated in this way should have no niche-DE genes, that is, 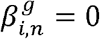 for all genes *g* and index-niche cell type configurations (*i,n*).

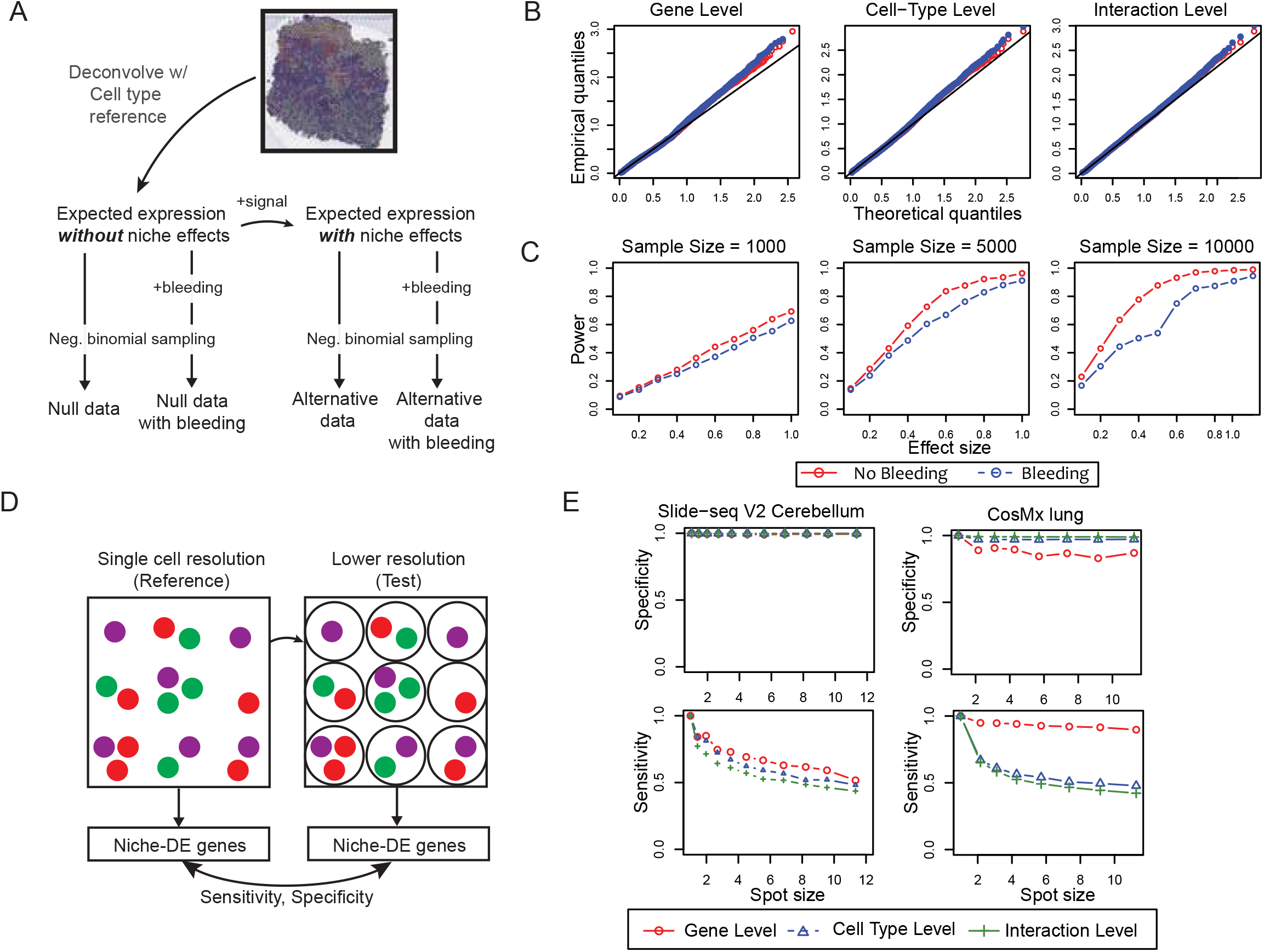
2A) Overview of data simulation: To simulate realistic ST data, we take a real ST dataset and perform deconvolution to calculate the expected expression vector for each spot *X_s_*. To generate data in the absence of niche effects, we simulate expression vectors from a negative binomial distribution with mean *X_s_* and overdispersion parameter 1. To generate data with niche effects, we specify *β_i,n_* for all index-niche pairs (*i, n*) and calculate the new expected expression vector for each spot *Y_s_* based on the niche-DE model. We then simulate expression vectors from a negative binomial distribution with mean *Y_s_* and overdispersion parameter 1. We also simulate ST data in the presence of spatial bleeding by calculating new expression vectors based on the SpotClean model with local bleeding parameter 0.25. 2B) Gene level, cell-type level, and interaction level null pvalue QQ plots when performing niche-DE on the simulated data. The empirical quantiles are based on those generated by niche-DE on the simulated data. The theoretical quantiles are based on the uniform distribution. 2C) Power calculation when performing niche-DE on the simulated data with niche-effects of varying size. 2D) Pseudo-spot data simulation overview: To simulate spot level data from single cell level data, we created pseudo spots by partitioning the field of view into equal-sized squares. Counts are aggregated within each square, to create a pseudo-spot. Spot size is defined as the average number of cells in a pseudo-spot. We applied niche-DE to these lower resolution datasets, and, using the niche-DE results from the original high-resolution datasets as the gold standard, we computed the sensitivity and specificity of the niche genes found at each spot size. 2E) Gene level, cell-type level, and interaction level Sensitivity and specificity vs spot size in both Slide-seq cerebellum and CosMX SMI NSCLC data.

Spot swapping is a known artifact in spatial transcriptomic data, where transcripts from a spot can “bleed” into nearby spots, contaminating the transcript pool of those nearby spots. This can lead to false spatial correlation and, as such, all ST datasets are contaminated to some degree. To examine how spot swapping may affect niche-DE p-values, we simulated spatial bleeding according to the SpotClean model where 25% of transcripts are bled into neighboring regions [8]. For each spot s, this results in a contaminated mean expression vector 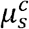. We then sampled a contaminated expression vector 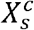 for the spot by drawing expression values from a negative binomial distribution with mean 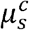. We denote by *X* = {*X_s_*} and 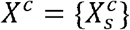 the simulated data sets with and without contamination, respectively.

To test the robustness of niche-DE to overdispersion and spot swapping, we perform niche-DE on *X* and *X^c^* at the gene, cell-type, and interaction levels. To visualize the p-values, we take their negative logarithm. In Fig. 2B, we see that the *quantile-quantile-plot* of niche-DE p-values on both *X* and *X^c^* are similar to that of a uniform distribution. This indicates that niche-DE effectively controls the type 1 error rate and is robust to spatial bleeding.

To examine if the power of niche-DE across effect sizes and dataset sizes is maintained under spot swapping, for 2000 random genes *G_β_*, we introduced a spike in effect 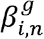 that varied within the set {0.2,0.4,0.6,0.8,1}. For *g* ∈ *G_β_*, let *μ_s,g,β_* be the expected expression of gene *g* in spot *s* according to the niche-DE model given 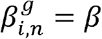. Similarly, let 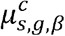 be the expected expression of gene *g* in spot *s* under the spot-swapping model with spike-in effect *β*. From *μ_s,g,β_* and 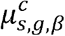 we can simulate *X_s,β_* and 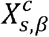 by drawing expression values based on a negative binomial distribution with mean *μ_s,g,β_* and 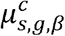 respectively. To simulate varying dataset sizes (number of spots), we bootstrapped *B* samples from the sets (*N_σ,s_, μ_s,β_*) and 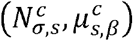 where *B* ∈ {1000,5000,10000}, *N_σ,s_* is the effective niche of spot *s* in the simulated dataset with no spatial bleeding, and 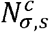 is the effective niche of spot *s* in the simulated dataset with spatial bleeding,. The results (Fig. 2C) show that there is minimal power loss due to spot swapping.

We believe that the robustness to bleeding is due to the bleeding effect being absorbed into the cell type proportion estimates during the deconvolution step. Suppose that cell type *i* in spot s bleeds a proportion *α* of its transcripts into neighbor spot *s*′. Perfect deconvolution based on marker genes would infer that spot *s*′ has an additional *α* cells of type *i*. If a gene *g* is a cell type *i* niche gene, it will be bled into neighboring spots more/less than what is expected, resulting in gene *g* being seen as a cell type *i* niche gene. Because neighboring cells have similar effective niches, this results in niche-DE detecting similar niche patterns in gene expression. Therefore, contaminated data will have similar niche patterns in gene expression to noncontaminated data. As such, the bleeding leads to biased estimates of cell type proportions, but the bias is not propagated to the niche-DE estimates.

### Recovery of niche-DE genes from ST data at varying spot resolutions

To quantify how much information is lost when using lower-resolution spot- and ROI-based technologies where each spot/region aggregates multiple cells, we simulated spot-level data by aggregating nearby measurements in two publicly available spatial transcriptomic data sets with sub-cellular resolution: the Slide-seq cerebellum data from [9] and the Nanostring CosMx SMI non-small cell lung cancer data from [16]. To simulate lower-resolution data, we created pseudo spots by partitioning the field of view into equal-sized squares (Fig. 2D). Counts are aggregated within each square. We call the average number of original spots/cells that are contained in each pseudo-spot the ‘spot size’. A larger spot size corresponds to a coarser dataset. We applied niche-DE to these lower resolution datasets, and, using the niche-DE results from the original high-resolution datasets as the gold standard, we computed the sensitivity and specificity of the niche genes found at each spot size (Fig. 2E).

For the Slide-seq cerebellum data, across all spot sizes, the specificity is maintained at almost exactly 1, indicating that type I error is effectively controlled at all spot resolutions. However, we see a clear trend of sensitivity decreasing as spot size increases. When the spot size is 4 cells, niche-DE recovers about 75% of the gold standard detections, and when the spot size increases to 10 cells, the recall rate drops to 50% at the gene, cell type, and interaction levels. This trend may be also be in part due to sample size decrease, since, when the spot size is 10, the corresponding coarser dataset has 10 times less data points.

For the CosMx SMI non-small cell lung cancer data, we see a similar trend: Across all spot sizes, specificity is maintained at almost exactly 1 for the cell type and interaction level tests, and above 90% for the gene-level tests. Sensitivity decreases, as expected, with increasing spot size. For this data set, recall rate at the gene level is maintained above 90% even at spot-size of 10 cells. However, as for the Slide-seq cerebellum data, recall rates for the cell-type- and interaction-level tests drop substantially as spot-size increases, leveling at 50% when the spot size is 10. Unlike the slide-seq data, the drop-off in sensitivity is sharper when spot-size increases from 1-2. We believe that this is due to the larger number of cell-types in this data set (20 versus 8), and thus the number of niche-index interactions tested across colocalizing cell types makes it hard to find exactly which index-niche pair(s) are driving the significance of a niche-DE gene.

These results indicate that, while there is indeed loss of power with decreased spatial resolution, over half the signals detectable at the sub-cellular resolution can be detected by niche-DE at a spot-size of 10 cells, without increasing false-positive rate.

### Inferring Ligand-Receptor Interactions via niche-DE

To determine which extracellular signaling mechanisms are driving niche-DE patterns between index cell type *i* and niche cell type n, we developed a procedure integrating Niche-differential genes, ligand expression, receptor expression, and Niche-Net [[11] data which links ligands with downstream target genes (Fig. 3A). The procedure, illustrated in Fig. 3B, starts with the ligandtarget matrix *A* = {*A_l,g_*: *l* = 1,…, *L; g* = 1,…, *G*} obtained from Niche-Net, where *L* is a set of ligands and *G* is a set of target genes. *A_l,g_* reflects the confidence that ligand *l* can regulate the downstream expression of gene *g*. Complementing this, for index cell type *i*, niche cell type *n*, and kernel bandwidth *σ*, Niche-DE provides a *G* dimensional vector of one-sided *t*-statistics *B_σ,i,n_* = {*B_σ,i,n,g_*: *g* = 1,…, *G*}, where *B_σ,i,n,g_* reflects whether or not gene *g* is an (*i, n*)^+^ niche gene operating at kernel bandwidth *σ*. For each ligand *l*, we first identify the top downstream genes using |*A_l,g_*} (Fig. 3B step 1), and then calculate a ligand activity score for each niche and index cell type pair (*i, n*), which measures the degree to which these downstream targets are found to be niche associated between *i* and *n* (Fig. 3B step 2). The null distribution of this activity score can be determined, and the ligands whose activity score pass the null p-value threshold are assumed to be the most likely ligands expressed by niche cell type n in its interaction with index cell type *i*. We call this set of ligands *C_i,n_*.

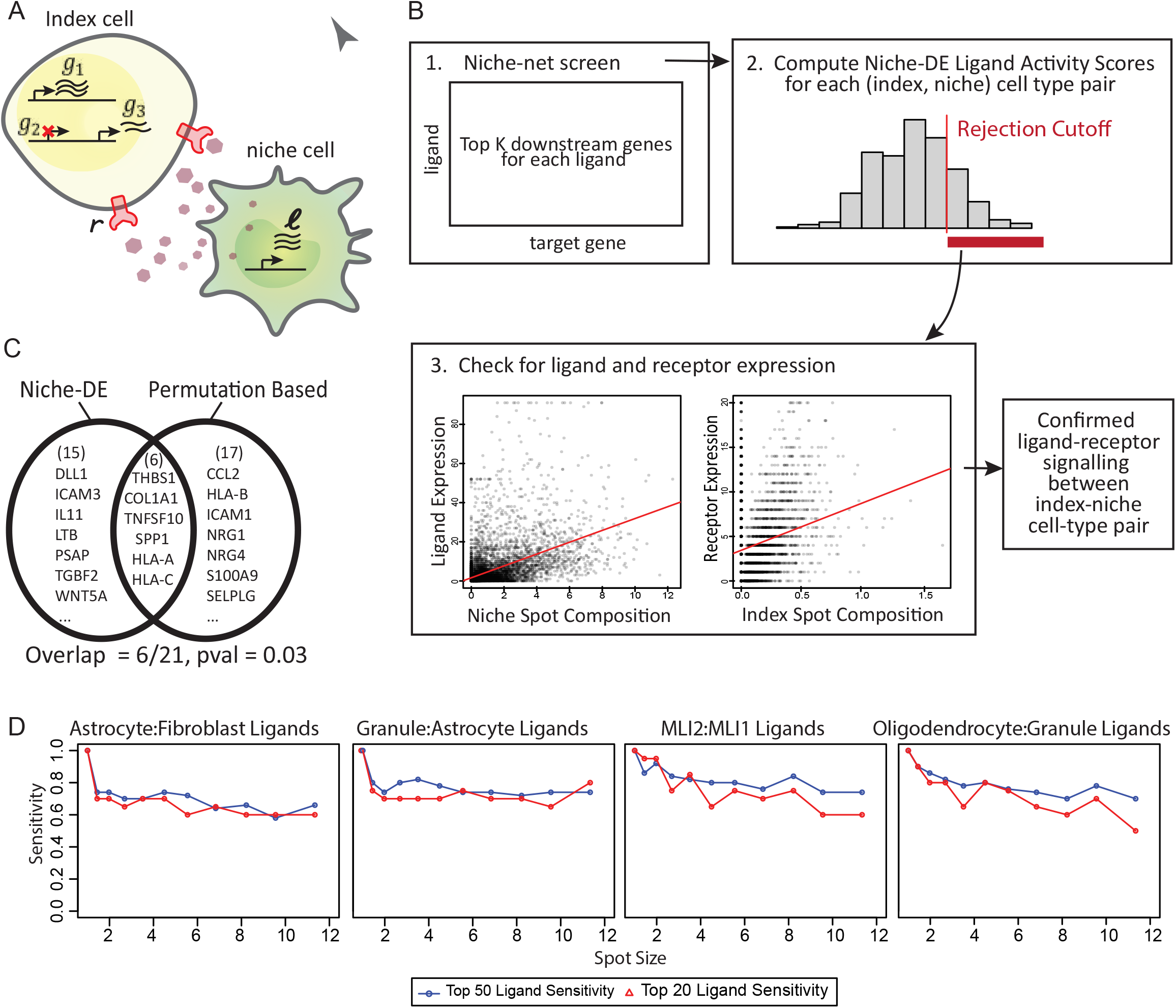
3A) Ligands from the niche cell type are received by the index cell type, resulting in a change in downstream gene expression. We expect these downstream genes to be (*index, niche*) genes. 3B) Overview of niche-DE Ligand-Receptor pipeline: We aim to determine which ligandreceptor pairs are active between the ligand expressing niche cell type *n* and the receptor expressing index cell type *i*. Using the ligand-target potential matrix from niche-net[11], we extract the top *K* downstream genes for each ligand. Using these downstream genes, we calculate a ligand activity score using the niche-DE T-statistic vector between index-niche pair (*i, n*) for the top *K* downstream genes. Ligands with an activity score greater than a threshold value are screened for expression in the niche cell type. 3C) To screen for ligand expression in the niche cell type we regress ligand gene expression on niche cell type spot composition. If the coefficient is significantly positive, we conclude that the ligand is indeed expressed by the niche cell type. We perform a similar regression to determine if the corresponding receptor is expressed by the index cell type. If a ligand and its corresponding receptor are found to be expressed by the niche and index cell types respectively, we conclude that the ligand-receptor is active between niche cell type *n* and index cell type *i*. 3D) Comparison of ligands inferred between the niche-DE based pipeline and a permutation based pipeline used by [16]. 3E) Sensitivity of top 20 and top 50 ligands by ligand activity score vs spot size using slide-seq cerebellum data from [5] as the reference.

Ligands in the candidate ligand set *C_i,n_* should be expressed by the niche cell type *n*, but this obvious condition has not yet been checked. Checking this condition is especially important since ligands may share similar downstream target genes, and thus, a ligand may have spuriously high activity scores due to lack of specificity in its Niche-net profile. Therefore, for each *l* ∈ *C_i,n_*, we perform a statistical test to confirm that the ligand is indeed expressed in the niche cell type *n*, and that at least one of its known receptors is indeed expressed in the index cell type *i* (Fig. 3B step 3). We call this combined approach niche-LR, for niche-ligand-receptor analysis. Details of niche-LR, including the computation of *C_i,n_* and the follow-up statistical test for niche expression of the ligand and index expression of the receptor, are described in Methods.

On the CosMx SMI NSCLC data, [16] proposed a permutation-based approach that assessed ligand-receptor co-expression between adjacent cell pairs. We compare the Niche-LR detected ligand-receptor pairs to those detected by the permutation-based method, see Supplementary Tables 6.1 and 6.2. Consider, for example, signaling between memory CD8 T (index cell type) and tumor (niche cell type). Out of the 21 unique ligands belonging to ligand-receptor pairs identified by Niche-LR, 6 ligands overlap with those identified by the permutation-based method (pvalue=0.03, Fig. 3C). The overlap (*THBS1, COL1A1, TNFSF10, SPP1, HLA-A, and HLA-C*) is significant, however the lack of overlap is also not surprising, given that the two approaches differ substantially: The permutation-based procedure, which is designed specifically for spatial transcriptomic data at single-cell resolution, ignores the expression of genes downstream of the ligand, and, by explicitly focusing on adjacent cells, ignores signaling between cells that are not immediate neighbors. Thus, genes downstream of the ligand does not need to show evidence of niche-DE for the ligand to be detected by [16]. However, [16] is expected to miss ligand-receptor channels that operate over larger distances, as well as those where RNA-level expression does not closely track the colocalization between the niche and index cell types.

To assess the accuracy of niche-LR at varying spot resolutions, we applied the method to the original and spot-aggregated datasets created as described in Figure 2D. The results (Fig. 3D) show that the top 20 and 50 candidate ligands in the Slide-seq cerebellum data are generally conserved across spot sizes and index-niche cell type configurations. In the CosMx SMI lung cancer data there is larger drop-off in overlap. We believe that this is due to the large number of cell types being screened (22 cell types leading to 484 index-niche configurations versus 8 cell types leading to 64 index-niche configurations for cerebellum), thus leading to higher numbers of colinear covariates in the regression and a vastly expanded number of parallel tests, decreasing the sensitivity for specific cell type configurations.

### Niche-DE detects tumor-fibroblast niche interactions with validation by CODEX imaging in liver metastases of colorectal carcinoma

We applied niche-DE and niche-LR to the integrative analysis of five 10X Visium samples obtained from liver metastases of colorectal carcinoma. For cell type deconvolution and mean expression profiles, we based our analysis on the reference scrna-seq data from [17]. Three of the samples (Patients 1-3) come from [18], while two additional samples (Patients 4-5) were generated in this study. CODEX data has been generated by [17] for patient 4. Supplementary Table 1.1 gives the details, including quality metrics, of these data sets. We deconvolved each sample using the single cell reference data set from [17], yielding the proportions of each of the 6 major cell types (hepatocytes, epithelial, fibroblast, macrophage, T lymphocytes, B lymphocytes) in each spot. Spatial maps of the estimated cell type proportions are shown in Supplementary Figure 1.2.

Since fibroblasts make up a substantial fraction of the cells in all samples, and since cancer-associated fibroblasts are an emerging target for anti-cancer therapy, we first focus on the niche interactions between fibroblasts and tumor cells. At FDR threshold of 0.05, niche-DE found 1226 genes whose expression in fibroblasts are significantly associated with the niche-enrichment of tumor cells. These genes are given in Supplementary Table 2.1. Pathway enrichment analysis of (*fibroblast, tumor*)+ genes identifies extracellular matrix organization, collagen production, and WNT signaling as the top three processes (Fig. 4B). This finding concurs with recent findings that collagen production and extracellular matrix remodeling are two of the most pronounced properties of tumor-associated fibroblasts versus their normal counterparts [17]. A full list of enriched pathways is given in Supplementary Table 2.2. A visual check of the spatial distribution for three of the niche-DE genes (*PSMD13, COL4A1*, and *COL15A1*), each in a separate patient sample, reveals that niche-DE gene expression are indeed enriched in regions of the tissue where fibroblasts and tumor cells co-localize (Fig. 4B). Such expression enrichment in regions of index-niche cell type colocalization is not always the case with niche-DE genes, as the niche-DE model uses flexible kernel bandwidths to allow for interactions beyond those between immediate neighbors.

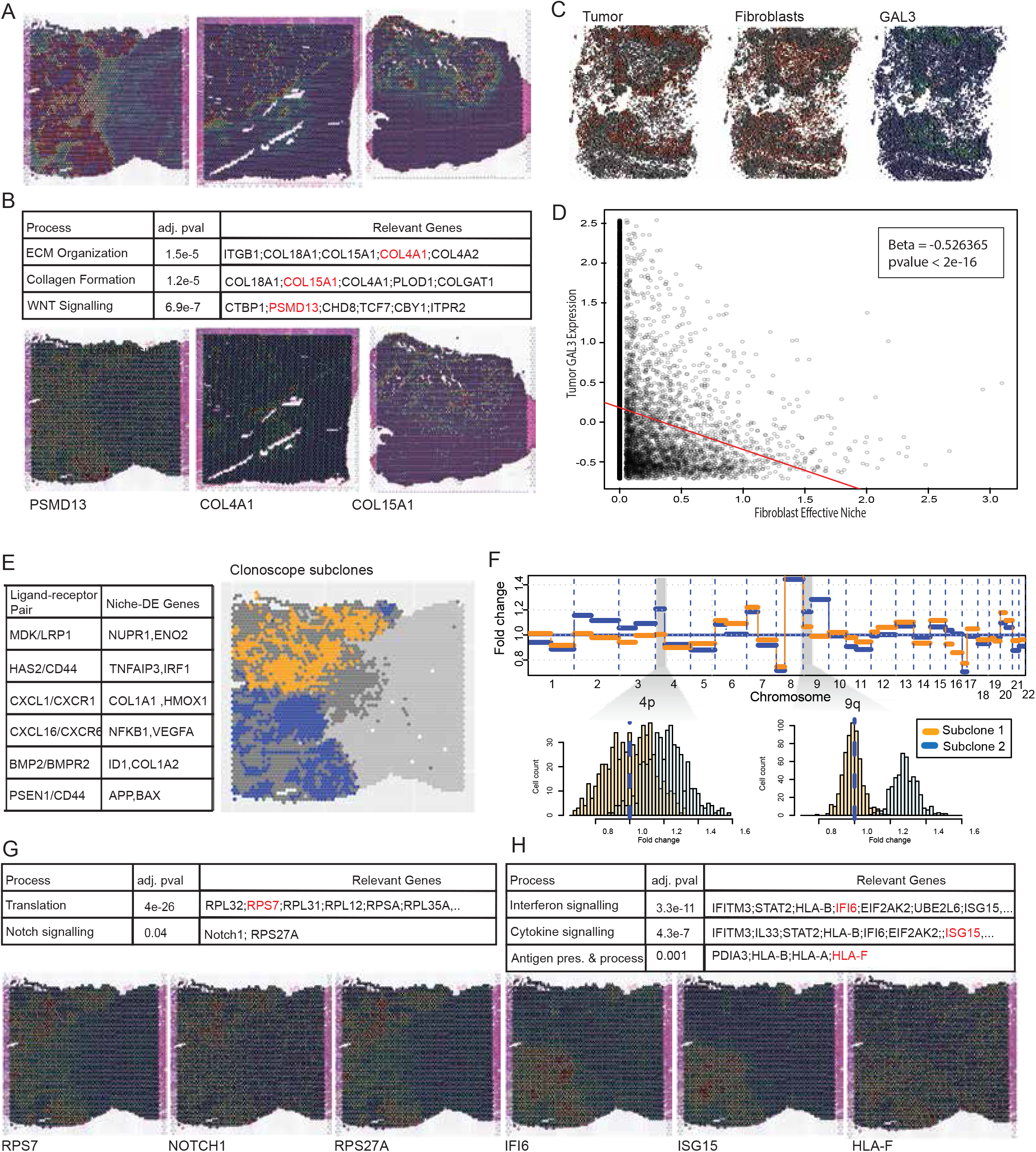
4A) Colocalization heatmaps between fibroblasts and tumor in liver samples 1,2, and 3. 4B) Using niche-DE marker genes in fibroblasts near tumor as input, pathway enrichment analysis finds ECM organization, collagen formation, and WNT signaling as top processes in fibroblasts in the presence of tumor cells. Spatial heatmaps of *PSMD13, COL4A1*, and *COL15A1* confirm expression of pathway related genes in the presence of tumor. 4C) Spatial heatmap of tumor abundance, fibroblast abundance, and GAL3 expression in CODEX data of liver metastasis of colorectal cancer in patient 4. 4D) Regression of tumor GAL3 expression on their fibroblast effective niche in CODEX data finds a significantly negative coefficient consistent with those found by niche-DE. 4E) Ligand-receptor pairs found between fibroblasts and tumor via niche-LR include *MDK/LRP1, HAS2/CD44*, and *CXCL1/CXCR1*. 4F) Clonalscope [20] finds two major tumor subclones in liver sample 1. Analysis finds an amplification of chromosomes 4p and 9q in subclone 2 relative to subclone 1. 4G) Using niche-DE marker genes in fibroblasts near subclone 1 relative to subclone 2 as input, pathway analysis finds translation and notch1 signaling and translation among enriched processes. Spatial heatmaps of *RPS7, NOTCH1*, and *RPS27A* confirm differential expression of pathway related genes in the region of the tissue containing tumor subclone 1. 4H) Using niche-DE marker genes in fibroblasts near subclone 2 relative to subclone 1 as input, pathway analysis finds interferon signaling, cytokine signaling, and antigen presentation among enriched processes. Spatial heatmaps of *IFI6,ISG15*, and *HLA-F*confirm differential expression of pathway related genes in the region of the tissue containing tumor subclone 2.

We also applied Niche-DE to identify genes whose expression in tumor cells are niche-associated with the enrichment of fibroblasts, with results shown in Supplementary Table 5.1. Among the top hits in patient 4 is *GAL3*, whose expression in tumor cells was found to be negatively associated with niche-enrichment of fibroblasts. Following-up on this finding, we analyzed the CODEX sample of patient 4 from [17]. The CODEX protein panel, which includes *GAL3*, is in Supplementary Table 5.2. Niche-DE analysis at the single cell level in CODEX detects a similar pattern of under expression of *GAL3* in tumor cells with increasing effective niche presence of fibroblasts (Fig.4C). A regression of tumor cell *GAL3* expression on the effective niche fibroblast concentration gives a slope of −0.53 (p-value<2^-16^, Fig. 4D).

We performed Ligand-Receptor analysis using niche-LR, detecting 26 unique ligands from tumor cells among the confirmed ligand-receptors between fibroblasts and tumor. This include Laminin family genes *LAMB2, LAMC2* which can bind to integrin family receptors on fibroblasts. Integrin signaling has been demonstrated to control fibroblast activation and response to mechanical signals in the TME [19]. Other ligand-receptor channels included *MDK/LRP1, CXCL1/CXCR1, HAS2/CD44*, and *BMP2/BMPR2* (Fig. 4E). The full list of ligand-receptor interactions identified from the data can be found in Supplementary Table 2.3.

### Niche-DE identifies subclone-specific niche interactions between tumor cells and fibroblasts

Tumor cells exhibit a large degree of heterogeneity due to subclonal evolution driven by both intrinsic genomic and epigenomic instability as well as extrinsic factors in its microenvironment. Thus, identifying different subclones of tumor cells in situ, and comparing the ways that the subclones interact with their local niche, may allow us to learn about mechanisms of tumor growth and invasion. Using Clonalscope [20], we identified 2 spatially distinct tumor subclones in Liver Sample 1. As shown in Figure 4F, subclone 1 is located predominantly at the upper left region of the slide, while subclone 2 is located predominantly at the lower left region of the slide (Fig. 4E). These two subclones are distinguished by chromosome 4p and 9q amplification in subclone 2 (Fig. 4F). Histopathological analysis of the liver sample shows that it is a conventional gland-forming adenocarcinoma. The region containing subclone 1 has less desmoplastic stroma, more architectural complexity, and some medullary-like features such as foci of a solid growth pattern compared to the region containing tumor subclone 2. Additionally, due to the more irregular and decreased gland formation, tumor subclone 1 is less differentiated relative to tumor subclone 2. Applying Niche-DE to this sample, with fibroblasts as the index cell type and each of the two subclones as niche cell types, we detect niche-differential expression that is specific to each subclone. Reactome analysis of marker genes of fibroblasts near subclone 2 reveal enrichment of interferon and cytokine signaling. Applying Niche-DE with subclone 2 as index cell type and fibroblasts as niche cell type identifies IFNGR1 to be a (*subclone* 2, *fibroblast*)^+^ niche gene as well. Reactome analysis of marker genes of fibroblasts near subclone 1 reveal enrichment of *NOTCH1* signaling and general translation to be enriched processes. These significant differences between fibroblasts that infiltrate distinct subclones at such close spatial proximity suggests that cancer-associated fibroblasts are highly plastic in adapting to subtle changes in their local tumor microenvironment. The full list of marker genes for each subclone can be found in Supplementary Table 4.1 and 4.2. The full list of enriched pathways associated with each index-niche cell type configuration can be found in Supplementary Tables 4.3 and 4.4.

### Niche-DE analysis identifies marker genes and signaling mechanisms specific to tumor associated macrophages in liver metastasis

We continue the integrative analysis of the Visium liver metastasis samples depicted in Figure 4 and Supplementary Figure 1.2. Reactome pathway analysis of (macrophage,tumor)+ genes identified by niche-DE revealed a significant enrichment of processes related to extracellular matrix (ECM) organization and metabolism [FIG 5A]. Ligand-receptor analysis between macrophages and tumor cells identify *CCL2/CCR2, CALR/LRP1*, and *IL18/CD48* as the ligand-receptor signaling mechanism driving these niche-DE genes [FIG 5B]. Full lists of confirmed ligand-receptor pairs, (macrophage,tumor)+ genes,and pathway analysis results can be found in Supplementary Tables 3.3 and 3.4, and 3.5.

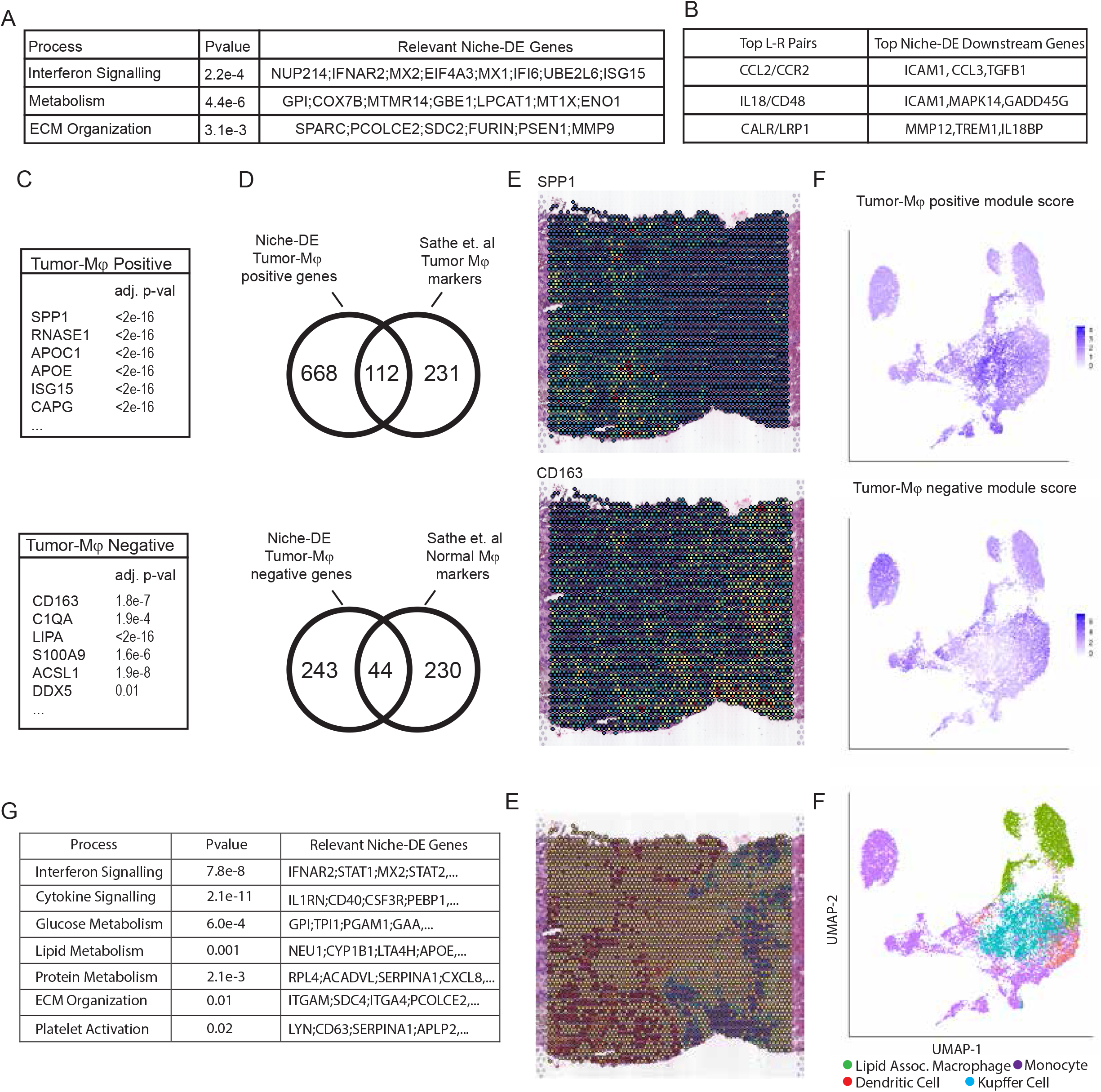
5A) Using niche-DE marker genes in macrophages near tumor as input, pathway enrichment analysis finds ECM organization, metabolism, and interferon signaling as top processes in macrophages in the presence of tumor cells. 5B) Ligand-receptor pairs found between macrophages and tumor via niche-LR include *CCL2/CCR2, IL18/CD48*, and *CALR/LRP1*. 5C) Lists of niche-DE positive and negative tumor associated macrophage marker genes. 5D) Overlap of niche-DE marker genes with lists found by Sathe et al. [17] 5E) From top to bottom. Top: Spatial heatmaps of *SPP1* and *CD163* confirm differential expression of marker genes in the region of the tissue containing tumor and hepatocytes respectively. Bottom: Differential spatial colocalization plots of macrophages near tumor and macrophages near hepatocytes. Larger values shown in red correspond to regions that contain macrophages near tumor and smaller values shown in blue correspond to regions that contain macrophages near hepatocytes. Regions in yellow correspond to regions with no macrophages or regions with macrophages that have a similar amount of tumor cells and hepatocytes in their effective niche. 5F) From top to bottom. Top: Featureplot of module scores for the set of niche-DE tumor-associated macrophage markers finds enrichment in the region of the UMAP corresponding to lipid-associated macrophages. Featureplot of module scores for the set of niche-DE normal macrophage markers finds depletion in the region of the UMAP corresponding to lipid-associated macrophages. Bottom: UMAP of scrna-seq from Wu et al. [18]. Cells were filtered to only include macrophages, monocytes, and dendritic cells. 5G) Using niche-DE marker genes in macrophages near tumor relative to hepatocytes as input, pathway analysis finds interferon signaling, cytokine signaling, and lipid metabolisms among enriched processes.

The liver microenvironment contains inflammatory macrophages derived from monocytes as well as non-inflammatory tissue resident macrophages such as Kupffer cells [21] (doi.org/10.1038/s41467-018-06318-7). Tumor-associated macrophages (TAM) represent a further reprogrammed macrophage subtype in the metastatic TME. Identifying and characterizing these cell states using only scRNA-seq is challenging given the similarity of their transcriptional profiles. However, in spatial transcriptomic data, we expect these two cell populations to be niche differentiable as macrophages from the normal liver should reside near hepatocytes while TAMs should be enriched near tumor cells. We developed an extension of the Niche-DE procedure to not only identify niche-DE genes, but to contrast the degree of niche-DE between two different niche configurations for the same index cell type. For example, we define TAM marker genes as genes whose expression in macrophages have significant positive association with tumor cell concentration in niche, but insignificant or negative association with hepatocyte concentration in niche. We define normal macrophage marker genes as the converse. Using this approach, at FDR threshold of 0.05 we identified 287 genes, including *HM0X1, CD163, LIPA, MS4A7*, and *CPVL*, as niche-DE marker genes of normal liver macrophages [FIG 5C]. These genes have been previously shown to be enriched in Kupffer cells from normal liver [21], thus validating this result. Niche-DE TAM marker genes included *TIMP2, APOE, SPP1*, and genes from the MMP family [FIG 5C]. 112 out of 780 niche-DE TAM markers genes also overlapped with previously identified scar-associated macrophages[22][FIG 5D]. The association of SPP1 upregulation with TAMs also confirms our previous finding, made using CODEX, of a *SPP1+* pro-fibrogenic TAM cell state in liver metastases [17], and the finding of a *SPP1+* TAM cell state in primary colorectal carcinoma [23]. Spatial heatmaps of *SPP1* and *CD163* confirm the localization of the expression of these genes in, respectively, the tumor and healthy regions of the tissue [FIG 5E]. Full lists of niche-DE TAM and normal macrophage marker genes can be found in Supplementary Tables 3.1 and 3.2.

To further follow-up on these results, we obtained single cell RNA-seq data [18] from paired samples of colorectal cancer and adjacent colon, liver metastasis and adjacent liver, lymph nodes along colon, and peripheral blood mononuclear cells (PBMC). On this data set, we computed module scores for the set of niche-DE tumor-associated macrophage markers and normal-associated macrophage markers identified above from the VISIUM data. We performed cell type classification via label transfer [12] from a liver cell atlas [24], and after subsequent denoising [25], compared the niche-DE module scores to the transferred cell type labels [FIG 5F]. For cells classified as lipid associated macrophages in the liver cell atlas, the niche-DE tumor-associated macrophage module score is high, while the niche-DE normal macrophage module score is low. In contrast, for cells labeled as Kupffer cells, the niche-DE normal macrophage module score is high, while the niche-DE tumor-associated macrophage module score is low. Reactome analysis of niche-DE tumor-associated macrophage marker genes finds glucose and lipid metabolism, as well as HDL mediated lipid transport as significantly enriched processes (Fig. 5G). A full list of enriched pathways can be found in Supplementary Table 3.6. These results are concordant with previous findings that metabolic reprogramming, and in particular lipid metabolic reprogramming, is a key factor in the regulation of TAM differentiation, polarization, and anti-tumor responses. [26].

## Discussion

We have presented a new method, niche-DE, for identifying cell type-specific genes whose expression is significantly up or down-regulated in the context of specific spatial niches. Niche-DE lends easily to integrative analysis, has robust type 1 error control, and allows for multiple kernel bandwidths for effective niche calculation. Via benchmarking studies conducted using single-cell and subcellular resolution spatial transcriptomic data, we showed that applying niche-DE to lower resolution spot-level data reliably recovers cell-cell interactions observed at the single cell level. The output of niche-DE can be used in a variety of ways, including but not limited to pathway enrichment analysis, ligand-receptor analysis, and niche marker gene analysis. In particular, we developed a procedure that integrates niche-DE statistics and niche-net ligand-target matrices to infer putative ligand-receptor signaling channels that underlie niche-differential gene expression. We illustrate the effectiveness of niche-DE through an integrative analysis of 10X Visium samples of liver colorectal cancer and corroborate our findings though parallel analyses of single cell RNA sequencing (scRNA-seq) and CODEX data for the same tissue type.

Niche-DE is based on a regression framework that regresses the observed gene expression at each cell/spot on the cell-type composition of its local neighborhood, which we call its effective niche. We start with a single-cell-level model where the parameters are easily interpretable, and from the single-cell-level model we derive a spot-level model with equivalent interpretation of parameters. By using negative binomial regression, we account for the presence of gene-specific overdispersion. Through a hierarchical false discovery rate control procedure, we achieve sensitivity for niche-DE genes under situations of high effective niche collinearity due to niche cell-type colocalization. Additionally, the niche-DE framework has a closed form expression for the joint distribution of the coefficients 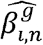, allowing for statistical inference to be done not only on the niche effects but also on the contrast between multiple niche effects. One application of such contrast testing is niche marker gene identification, which we perform on the liver metastasis data to delineate tumor-associated versus normal macrophage markers. The markers we identify corroborate with published findings on macrophage subtypes in liver metastases [[17] and primary colorectal carcinoma [[27], not only at the gene level but also at the level of cellular processes inferred through a Reactome analysis.

We also illustrate how niche-DE, when applied to tumor samples, enables the identification of subclone-specific interactions with local microenvironment. Niche-DE and subsequent Reactome analysis of fibroblasts within two tumor subclones in the liver metastasis data leads to the discovery of subclone-specific fibroblast subtypes: One subtype is enriched for interferon-stimulated gene signatures, while the other subtype is notably devoid of such signaling and is instead enriched for *NOTCH1* signaling. The juxtaposition of two different fibroblast subtypes interacting with different tumor subclones within such close spatial proximity highlights the functional heterogeneity of cancer-associated fibroblasts.

Niche-DE genes are assumed to be caused by local cell-cell communication, such as ligand-receptor signaling. We develop a method to infer which ligand receptor pairs are active based on niche-DE statistics. This allows the identification of receptor-ligand pairs spatially restricted to a localized niche, which indicate paracrine communication between cells. We show that while there is significant overlap between the ligand-receptor pairs found via our method and those found via permutation-based methods used in [16], both methods identify unique ligand-receptor pairs.

A limitation of niche-DE is that it only considers two-way interactions, e.g differential expression of macrophages genes near fibroblasts. This may be too simplistic for spatial transcriptomic datasets where two colocalizing cell types can be found in multiple global environments, e.g. macrophages near fibroblasts in tumor regions and macrophages near fibroblasts in healthy regions. One way to allow for interactions of more than 2 cell types is to augment the effective niche vector in niche-DE via the inclusion of all pairwise products between cell types. However, the number of hypotheses increases exponentially with the order of interaction, and although feasible, we believe the current data size and quality only allow sensitive detection of pair-wise interactions.

## Methods

### Derivation of the Spot-level Niche-DE Model

Model (1) specifies a log-linear dependence of expected gene expression on niche-composition. If the dependence were linear, then we could simply sum across all cells within each spot to derive an equivalent spot-level model with the same interpretations for the parameters 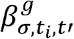. However, due to the gross variance inflation in gene expression data, it is necessary for the regression to be in log space. Here, we first show that the mean relationship in the single celllevel model can be approximated by a linear model, and then use that approximation to derive the equivalent spot-level regression formula. Starting with equation (1),

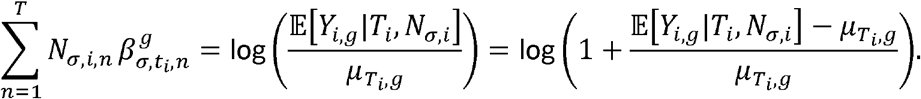

Using the approximation log(1 + *x*) ≈ *x* for *x* → 0, and assuming that the overall effect size 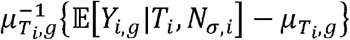 is small, the above leads to the approximation

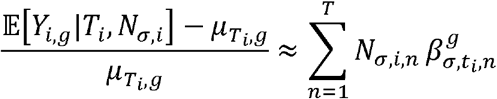

and thus,

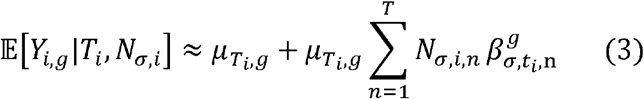

Hence, the cell-level loglinear model can be approximated, at the mean level, by a linear model, but with the extra *μ_T_i_,g_* factor multiplied to the effect size for each cell.

Now consider spot-level data. Let *s*(*i*) denote the spot where cell *i* resides and *X_s,g_* be the observed expression of gene *g* in spot *s*. Using the same kernel function, we compute an effective niche for each spot *s*, which approximates the effective niche for each cell *i* where *s*(*i*) = *s*. Since spots can contain a multitude of cell types, let *n_s_* be the cell-type composition vector for spot s, i.e. *n_s_* = (*n_s,t_*: *t* = 1,…*T*), where *n_s,t_* is the number of cells of type *t* in spot *s*. We sum the approximations (3) across all cells within each spot to yield

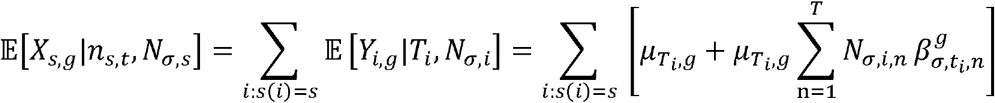

Because all single cells in the same spot have the same effective niche and the sum only depends on the cell type of each cell, by summing over the niche cell type rather than each single cell we get an alternative sum of

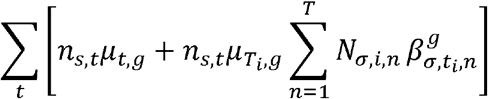

Let *μ_s,g_* = ∑^t^ *μ^t,g^* and 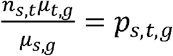. The sum above can be simplified to

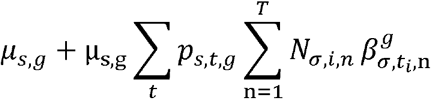

Moving terms in the sum we get that

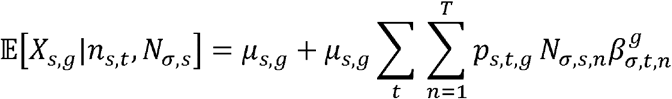

Subtracting *μ_s,g_* and then dividing by *μ_s,g_* on both sides of above, we have

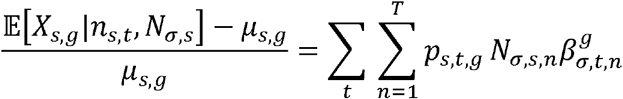

We expect the left-hand-side of the above to be small, and thus, using the same *x* = log(1 + *x*) approximation, we get our final spot level model for expected expression of gene *g* in each spot conditioned on the spot’s cell type composition and effective niche,

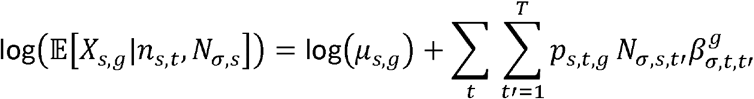

Note that, since 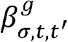 is the same parameter carried through from model (1), it has the same interpretation as model (1), that is, it represents the effect of unit niche-composition increase of cell type *t*’ on the single-cell expression of gene *g* in cell type *t*.

### Parameter estimation in the Niche-DE model

#### Data Pre-processing

To remove outliers in gene expression, for each gene, we cap gene expression to the 99.5 percentile across all spots.

#### Deconvolution and estimation of *n_s,t_*

All samples used were deconvolved with RCTD using the “Multi” setting with maximum number of cell types in each spot equal to 4. We thresholded each deconvolution estimate to have a minimum value of 0.05. Because we found that deconvolution via RCTD overestimated the amount of tumor cells and hepatocytes in each spot, we thresholded each deconvolution estimate such that tumor cell and hepatocyte composition had a minimum value of 0.25. Spots that did not meet this criterion had tumor cell and hepatocyte composition set to 0.

Critical to the effective niche calculation is an estimate of the number of cells in each spot *n_s,t_*. Assume that there are *N_s_* cells in spot *s*, for which the cell type composition is *π_s,t_*. From the reference dataset, let *L_t_* be the average library size of cell type *t*. The expected library size for spot *s* is given by *N_s_*. ∑*_t_π_s,t_ L_t_*. Let *L_s_* be the observed library size of spot *s*. A natural estimate for *N_s_* given 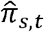, the estimate of the cell type composition of spot *s*, is thus 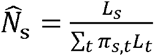. Using this estimate, we can then approximate *n_s,t_* by 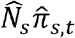.

#### Joint analysis of multiple spatial transcriptomic data sets of different resolution

Because niche-DE only depends on the effective niche, joint analysis of multiple spatial transcriptomic data sets, possibly of different resolutions, can be accomplished easily by ensuring that the effective niche is on the same scale across all data sets. We will demonstrate how to do this for two data sets, the first we call the reference dataset and the second we call the query dataset. The first step is to scale the coordinates of the query dataset such that one pixel in the query dataset corresponds to same physical distance as one pixel in the reference dataset. This makes sure the kernel bandwidth used has the same interpretation across both datasets.

If both datasets are of spot-level resolution, as is the case with 2 Visium datasets, we must scale the effective niche of the query dataset as well. The reason is that the estimated number of cells in each spot is determined by the library size of the spot. Since the niche-DE model is a regression on the effective niche, the effective niches of the query and reference dataset must be corrected to account for sequencing depth. To accomplish this, let the spot radius of the reference and query datasets be *r* and *q* respectively. Let *N_r_* and *N_q_* be the average number of cells in each spot of the reference and query datasets. We propose scale the effective niche of the query dataset by (*N_r_q*^2^)/(*N_q_r*^2^). Scaling by *N_r_*/*N_q_* corrects for differences in sequencing depth between the reference and query dataset. Scaling by *q*^2^/*r*^2^ makes sure that our correction is not driven by differences in spot sizes in the two datasets. For example, if a spot doubles in size, it will occupy 4 times the area and thus its library size is expected to be 4 times higher.

#### Normalization and Filtering

In single cell data, cells of the same type can have different library sizes. As such we must adjust *μ_i,g_*, the expected expression of gene *g* for single cell *i*, to account for this before applying niche-DE. Let *L_T_i__* be the expected library size of a cell of type *T_i_*. Let *L_i_* be the library size of single cell *i*. We scale *μ_i,g_* to be such that *μ_i,g_* = *A_i,g_L_i_*/*L_T_i__*. w, where *A_i,g_* is the average expression of gene *g* in cell type *T_i_* as given by the reference. This ensures that niche genes are not driven by patterns in sequencing depth. Note that in spot level data, this scaling is a by product of the estimation of *N_s_* and its role in calculating *μ_s,g_* the expected expression of gene g for spot *s*.

Let *N* be the effective niche matrix of our dataset where *S* is the number of spots and *T* is the number of cell types. Before running the niche-DE model, we scale *N* column wise so that the column mean of non-zero entries is 1. This alleviates biases in overall tissue cell type composition so that the *β* coefficients are on the same scale. We then center the columns to have mean 0. We also set *p_s,t,g_* = 0 if it is less than 0.05. This was shown to increase numerical stability.

Also for the sake of numerical stability, we set 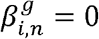 unless three conditions hold. The first is that the total expression of gene *g* across all spots is greater than some threshold *C*. The second is that there exists more than *M* spots that contain index cell type *i* which have an effective niche that contains niche cell type n. This is to ensure that signals found are not dominated by very few points. The third condition is that gene *g* is in the top *γ* percentile of expressed genes in cell type *i*. This is because most genes should have essentially 0 expression in a given cell type *i*. Default values for *M*, and *γ* are 30 and 80 respectively. *C* is dependent on the number of spots in the dataset but niche-DE is quite robust to the choice of *C*.

Finally, for computational efficiency, rather than using standard Negative binomial regression which is quite slow, we instead perform standard Poisson regression using the “glm” package in R. Then with the given regression coefficients, we estimate the overdispersion parameter with the “optim” function in R. This was shown to work well as seen in figure 2A.

#### Parameter Choice

Recall that the niche-DE has three parameters: *C*, the minimum expression across all spots necessary for a gene to be included in the regression, *M*, the minimum number of spots that contain the index cell type and have the niche cell type in its effective niche for an index-niche pair to be included in the regression, and *γ*, the minimum percentile a gene’s expression needs to attain in the index cell type to be included in the regression. For the integrated 10X VISIUM colorectal cancer data, we set (*C, M, γ*) = (400,10,80%). For the SMI NSCLC data, we set (*C, M, γ*) = (6000,50,60%). Forthe Slide-seq cerebellum data, we set (*C, M, γ*) = (30,30,80%).

Niche-DE on the COSMX data was done with kernel bandwidths of 150,250, 350, and 450 pixels which correspond to 30, 50, 70, and 90 *μ*m respectively. Niche-DE on the slide-seq data was done with kernel bandwidths equal to the first, fifth, and tenth percentile of the total distance matrix of our dataset. Niche-DE on the integrated data was done with kernel bandwidths 1,200, and 500 which corresponded to bandwidths of 0.2,100, and 250*μ*m.

### Downstream Analysis

#### Reactome Analysis

Let *G_i,n_* be the set of genes that are found to be (*i, n*)^+^ niche genes. To infer what processes are active in index cell type *i* in the presence of niche cell type *n*, we use “enrichR” [[28]] to perform a pathway analysis using *G_i,n_* as input. The database we use is “Reactome 2016”.

#### Marker Gene Analysis

Given index cell type *i* and niche cell types *n*_1_, *n*_2_, we define a gene *g* to be a (*i, n*_1_) marker if the degree of upregulation in cell type *i* in the presence of cell type *n*_1_ is significantly greater than the degree of niche upregulation in the presence of cell type *n*_2_. Statistically, this is equivalent to performing a contrast test with null hypothesis 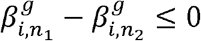 against alternative hypothesis 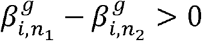. Because niche-DE is based on a negative Binomial regression, there is a closed form expression for the joint distribution of *β*. The distribution of 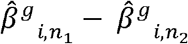 can be shown to be normally distributed with a mean of 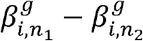 and a variance that can be computed. We perform marker gene analysis to find marker genes in macrophages near tumor and hepatocytes on an integrated dataset containing 4 liver metastasized colorectal carcinoma Visium datasets.

#### Module Score Computation, Labelling and Denoising of ScRNA-seq Data

To evaluate the validity of our niche-DE tumor associated macrophage marker gene set, we calculate the module score of this set of genes on liver CRC scRNA-seq data from Wu et al [[20] using the “AddModuleScore” function in Seurat applied to SAVER [25] denoised scRNA-seq values. Because the scRNA-seq data is not labelled, we perform label transfer using Seurat with the cells from the liver cell atlas [[29] as the reference. Cells that were classified to be myeloid were extracted and label transfer was performed again using myeloid subtypes from the liver cell atlas as reference.

#### Ligand-Receptor Analysis

To determine which extracellular signaling mechanisms are driving niche-DE patterns between index cell type *i* and niche cell type *n*, we developed a procedure integrating Niche-net [11] and Niche-DE statistics. The two statistics reflect different types of evidence supporting intra-cellular signaling: Niche-net provides a ligand-target matrix *A* = {*A_l,g_: l* = 1,…, *L; g* = 1,…, *G*], where *L* is a set of ligands and *G* is a set of target genes. *A_l,g_* reflects the confidence that ligand *I* can regulate the downstream expression of gene *g*. For index cell type *i*, niche cell type *n*, and kernel bandwidth *σ*, Niche-DE provides a *G* dimensional vector of one-sided *t-*statistics *B_σ,i,n_* = (*B_σ,i,n,g_: g* = 1,…, *G*], where *B_σ,i,n,g_* reflects whether or not gene *g* is an (*i,n*)^+^ niche gene operating at kernel bandwidth *σ*.

Studies suggest that different ligand tend to be effective over different distances [30]. To infer if cells of type n within the niche of index cells of type *i* are signaling to the index cells via ligand *l*, we first compute the optimal kernel bandwidth 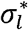 for ligand *l* based on the niche-DE regression likelihood score. This represents the most likely bandwidth at which we expect the ligand to operate and, by extension, the bandwidth at which we expect to observe downstream niche-DE genes. We then extract the top *K* downstream target genes of ligand *ℓ* from the Niche-net ligand-target matrix, which we call *g*_1_,… *g*_K_. Using the Niche-Net ligand-target matrix values, we compute a weight vector *W*(*l*), where 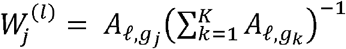.

Then, we combine these with the Niche-DE statistics to compute a ligand activity score 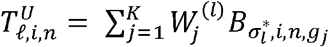. If the top downstream genes {*g*_1_,…, *g_K_*] are not niche-DE in index-niche configuration (*i, n*), then 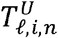 should abide by the null distribution and thus be approximately normally distributed with mean 0 and variance 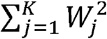. As such, we compute the standardize ligand activity scores, standardizing 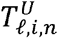 by its standard error under the null,

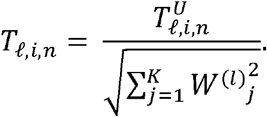

Thus, between index cell type *i* and niche cell type n, we compute *T_l,i,n_* across all ligands *l*. We sort them in decreasing order, and let *T_(M),i,n_* be the *M*^th^ largest order statistic amongst all ligand activity scores between index cell type *i* and niche cell type n. We define the candidate ligand set as *C_i,n_* = {*ℓ*:*T_ℓ,i,n_* > max(1.64, *T_(M),i,n_*)}. The candidate ligand set represents the ligands with top evidence for being involved in signaling between index cell type *i* and niche-cell type *n*, based on the Niche-Net matrix and the niche-DE gene expression patterns between the two cell types.

Ligands in the candidate ligand set *C_i,n_* should be expressed by the niche cell type *n*, but this obvious condition is not a pre-requisite for its selection and still needs to be checked. This check is especially important since ligands may share similar downstream target genes, and thus, a ligand may have spurious high activity scores due to lack of specificity in its Niche-net profile. Therefore, we filter ligands and their receptors to ensure that the ligands are indeed expressed in the niche cell type *n*, and that the receptors are indeed expressed in the index cell type *i*, for (*i, n*) pairs where ligand *I* ∈ *C_i,n_*. This filter operates as follows: If a candidate ligand *ℓ* is indeed expressed by cell type *n* in the vicinity of index cell type *i*, there should be a positive correlation between the count of ligand *ℓ* in a spot and the abundance of cell type *n* in the spot, for spots that have index cell type *i* in their effective niche. Thus, we perform a poisson regression of the observed ligand expression *X_s,ℓ_* on the inferred spot-by-cell-type composition matrix (*n_s,t_*), yielding coefficient vector 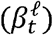 for the enrichment of ligand *I*within each cell type *t*. In this regression, we filter to only include those spots which have index cell type *i* in their effective niche evaluated at bandwidth 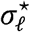. From the regression, we get a pvalue for testing 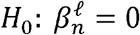 versus 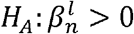, which we call 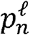. After applying the BH procedure over all candidate ligand pvalues 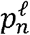 for ligands in *C_i,n_* at a pre-set false discovery rate *α*, we conclude that those with significant adjusted pvalues are indeed expressed in niche cell type *n* around index cell type *i*, and is reported as the set of “confirmed ligands” 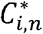. If the spatial data were single cell resolution, we instead determine whether ligand *ℓ* is in the top *α*% of genes expressed by niche cell type *n* that have cell type *i* in their effective niche evaluated at bandwidth 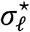, with *α* specified by the user.

Using the ligand receptor list from [31], we can furthermore curate a list of confirmed receptors for each ligand. Similar to above, to determine whether index cell *i* expresses the corresponding receptor *r_l_* to confirmed ligand 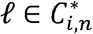, we perform a poisson regression of the observed receptor expression *X_s,r_l__* on the inferred spot composition matrix (*n_s,t_*) yielding coefficient vector 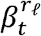. For this regression, we only include those spots which have niche cell type *n* in their effective niche evaluated at bandwidth 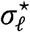. This is to ensure that we only consider spots that can potentially receive ligand *ℓ* emitted by niche cell type *n*. After computing pvalues for 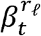 and performing the BH procedure over all known receptors of ligands in 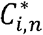, we conclude that those receptors with significant adjusted p-values are indeed expressed in index cell type *i* in the vicinity of niche cell type *n*. If the spatial data were single cell resolution, we instead determine whether receptor *r_ℓ_* is in the top *α*% of genes expressed by index cell type *i* that have cell type *n* in their effective niche evaluated at bandwidth 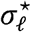. If ligand *ℓ* and its corresponding receptor *r_ℓ_* are confirmed to be expressed in the niche and index cell types, respectively, we then conclude that the ligand receptor pair (*ℓ,r_ℓ_*) is active in signaling between index cell type *i* and niche cell type *n*.

The user tunable parameters of this process are *K*, the number of downstream genes to consider from the Niche-Net matrix, *M*, the number of candidates to include in the candidate ligand set, and *α*, which is either the false discovery rate cutoff (for spot-level spatial data) or the expression rank cut off (for single cell resolution spatial data). We performed ligand-receptor analysis on the CosMx SMI NSCLC data as well as the integrated set of 10X Visium colorectal cancer datasets. We use (*K, M*) = (50,50,0.05) for the SMI data and (*K, M*) = (25,50,0.5) for the Visium data.

### Simulation Designs

#### Bleeding Simulations

##### Generic Model

First, we describe how to simulate from the null model of no niche-effects, where the gene expression of a spot/cell depends only on its cell type composition/identity. Given a ST dataset, let *n_s,t_* be the estimated cell type composition vector for spot *s*, i.e. *n_s,t_* is the proportion of cells at spot *s* that are of cell type *t. n_s,t_* can be found via deconvolution [[9]]. If the data were single-cell resolution, then *n_s,t_* is simply a cell type label, *n_s,t_* = 1 if cell s is assigned label *t*, and *n_s,t_* = 0 otherwise. We multiply *n_s,t_* with the reference cell-type mean gene expression matrix to give us the expected expression vector *μ_s_* = (*μ*_*s*1_…, *μ_sG_*), where *μ_sg_* is the expected expression of gene *g* at spot/cell *s* without any niche effects. After library size normalization of *μ_s_* we can sample an expression vector *X_s_* by drawing expression values based on a negative binomial distribution with mean *μ_s_* and gene specific overdispersion vector *γ_g_*. The coordinates of simulated spot *s* will be the same as the real coordinates of spot *s*. This can be done with any spatial transcriptomics data, but for our simulations, we use Visium from patient 4 (referenced below) and set *γ_g_* = 1 for all genes *g*.

##### Generic Model Incorporating Niche Effects

Let the assumed underlying spatial kernel bandwidth for gene *g* be *σ*. Let *N_σ,s_* be the effective niche vector for spot s after appropriate normalization. Letting 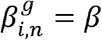 and 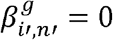 for all *i*′ ≠ *i* and *n*′ ≠ *n*, we have that

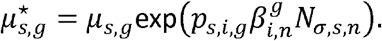

Thus we sample *X_s,g_* by drawing expression values based on a negative binomial distribution with mean 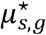 and gene specific overdispersion vector *γ_g_*. To simulate datasets of size *B*, where *B* is larger than the total number of cells/spots in the initial real data, we bootstrap *B* samples from the set of (*N_σ,s_*, 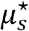) For each spot *s*′ that is sampled, we simulate a spot with an expression vector 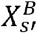 sampled from a negative binomial regression with mean 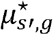 and an effective niche equal to *N_σ,s′_*. To test the power of niche-DE across varying effect sizes and sample sizes, we vary 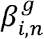 within {0.1,0.2,…,1} and vary *B* within {1000,5000,10000}. For our simulations, we use patient 4 (referenced below). We set *i* to be fibroblasts, *n* to be tumor, and *σ* to be the first percentile of the total distance matrix of our dataset. We select 2000 random genes to be (*i, n*) + niche genes.

##### Spatial Bleeding Model

Spot swapping is a known artifact in some spatial transcriptomic data sets where transcripts “bleed” into nearby spots, inducing artifactual correlation between the transcript counts in adjacent spots [REF]. The severity of spot swapping varies across data sets. To examine how bleeding may affect niche-DE analysis, we add the bleeding effect to our simulation data according to SpotClean’s model [REF]. In particular we let *α_g_* be the gene specific local bleeding parameter. Let *K_τ_* be a gaussian kernel with bandwidth *τ*. Letting *μ_s,g_* be the underlying expected expression of gene *g* in spot s with no bleeding effect and 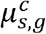 be the expected expression of gene *g* in spot *s* in the presence of bleeding, under the SpotClean model,

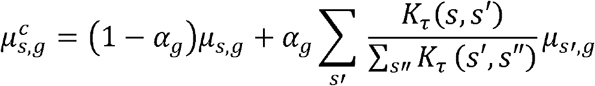

We then sample the observed gene expression from a Negative Binomial with mean 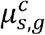 and gene-specific dispersion. We introduce this spatial bleeding effect to both the generic null model and the generic model with niche-effects. This allows us to observe the effect of bleeding on both null pvalues and the power of niche-DE. We set *τ* equal to the first percentile of the total distance matrix of our dataset.

Note that, with the spot-swapped simulation data, the procedure starts with a deconvolution which yields a contaminated estimate of spot cell type composition 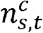. This contaminated estimate 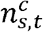 is then used to generate a contaminated effective niche vector for each spot. Thus, the spot-swap effect is “absorbed” in the cell type composition estimation, which alleviates the influence on downstream niche-DE analysis.

#### Pseudo-Spot Data Generation and Specificity and Sensitivity Analysis

##### Generating Pseudo-spot Data

To simulate lower resolution datasets, we create pseudo spots by partitioning the field of view of a reference dataset into squares of side length r. All cells in the same square are assigned to the same spot. Gene expression in each spot is calculated by aggregating the gene expression of all cells in the spot. We create pseudo-spot data using the COSMX SMI NSCLC and Slide-seq cerebellum data as the initial datasets. The radii used for the COSMX data were 100, 125, 150, 175, 200, 225, and 250 pixels which corresponds to 20,25,30,35,40,45, and 50*μ*m, respectively. The radii used for the slide-seq data were 10,25,30,35,40,45,50,55,60,65,70, and 75 pixels.

##### Sensitivity and Specificity Calculations

Performing niche-DE on the initial high-resolution dataset, we obtain a set of significant (*i, n*) niche genes at the gene, cell type, and interaction levels. Using this set of genes as a reference, we can perform niche-DE on the pseudo-spot dataset as well to calculate the sensitivity and specificity at all three levels.

Niche-DE on the COSMX SMI NSCLC pseudo-spot data was done with kernel bandwidths of 150, 250, 350, and 450 pixels which correspond to 30, 50, 70, and 90*μ*m respectively. Niche-DE on the slide-seq pseudo-spot data was done with kernel bandwidths equal to the first, fifth, and tenth percentile of the total distance matrix of our dataset.

##### Ligand Sensitivity Calculations

To test the robustness of our ligand-receptor inference method, we apply the method to obtain the top *M* ligands by activity score for the reference COSMX and slide-seq datasets as well as the corresponding pseudo-spot datasets. The reason we chose to compare ligand-activity scores rather than candidate ligands or confirmed ligand-receptor pairs was because niche-DE applied to the lower resolution datasets has less power than when it is applied to the original datasets. This makes the ligand potential scores lower which reduces the number of candidate ligands. As a result, the comparison of candidate ligand sets much noisier. By comparison, the set of top *M* ligands is more robust as we still expect the ligand potential scores across resolutions to have a similar order. We choose *K* = 50 and *M* ∈ {20,50}.

### Datasets and Data Analysis

#### Published data sets

- **Slide-seq cerrebellum:** We use the pre-deconvolved slide-seq cerebellum data from [9]. This data can be found at https://singlecell.broadinstitute.org/singlecell/study/SCP948 and is the file named ‘myRCTD_cerebellum_slideseq.rds’. This dataset contained 19 cell types, 11626 spots of of spatial resolution 10*μ*m, and 5034 genes. For all analyses, we filtered the estimated cell type composition vectors to only include the top 8 most ubiquitous cell types. This corresponded to Astrocytes, Bergmann cells, Fibroblasts, Granule cells, MLI1, MLI2, Oligodendrocytes, and Purkinje cells.
- **COSMX SMI Non-small Cell Lung Carcinoma:** We obtain Nanostring CosMx SMI non-small cell lung cancer data from [16]. We used Lung sample 9 which contains two samples measuring 960 genes with 87684 and 139735 cells and 22 cell types. Each sample spans a 5mm by 4.25mm area. Niche-DE and downstream analyses were performed on both samples in an integrative fashion. Permutation based ligand-receptor analysis was done only on the sample with 139735 cells.
- **10X Visium (patient 4):** This data contains liver metastasis of colorectal cancer. It has 848 spots and 36601 genes.
- **10X Visium (patient 5):** This data contains liver metastasis of colorectal cancer. It has 1663 spots and 36601 genes.
- **CODEX (patient 4):** CODEX data of liver metastasized colorectal cancer from patient 4 was obtained from [17]. It contains 33812 cells and 25 markers.
- **Wu et al. 10X Visium (Patients 1,2,3):** We obtain 3 10X Visium data sets from liver metastasized colorectal cancer corresponding to patients 1,2,and 4 of [18]. These datasets contain 3826, 4658, and 3721 spots with 36601 genes. The dataset corresponding to patient 1 was found to have two tumor subclones found by Clonalscope.
- **Wu et al. scRNA-seq:** We obtain single cell RNA-seq data from paired samples of colorectal cancer, adjacent colon, liver metastasis, and adjacent liver, lymph nodes along colons, and peripheral blood mononuclear cells (PBMC) from [18]. After filtering the data to only include macrophage/monocytes as explained in section 3.3, the final dataset contains 17791 cells.
- **Colorectal cancer and liver metastasized colorectal cancer scRNA-seq:** For all 10X Visium colorectal cancer datasets, we merge scRNA-seq datasets from [17]. The cell types present are Hepatocytes and NK cells, Cholangiocytes, T cells, B/Plasma Cells, Tumor, Endothelial, Macrophages, and Fibroblasts. Because all samples contained little to no lymphocytes and endothelial cells, we filter all deconvolution results to only include Hepatocytes and Cholangiocytes, Tumor, Macrophages, and Fibroblasts.

#### Sample acquisition For Visium Samples 4 and 5

This study was conducted in compliance with the Helsinki Declaration. Patients were enrolled according to a study protocol approved by the Stanford University School of Medicine Institutional Review Board (IRB-44036). Written informed consent was obtained from all patients. Samples were surgical tumor resections.

#### Tissue processing

Tissues were placed in cryomolds containing chilled TissueTek O.C.T. Compound (VWR). Additional O.C.T. was added to cover the tissue and cryomold was placed on powdered dry ice. Blocks were sealed and stored at −80°C.

#### 10× Visium Spatial transcriptomics library preparation

Libraries were prepared using Visium Spatial Gene Expression Reagent Kit (version 1) (10X Genomics). All steps were performed according to the manufacturer’s protocol. Briefly, 10 μm thick tissue sections were placed onto a Visium Spatial Gene Expression slide using a cryostat. Following methanol fixation, hematoxylin and eosin staining was performed, coverslip was mounted, and slides were imaged using a Leica DMI 6000 or Keyence BZ-X microscope. A permeabilization time of 18 minutes was used, which was determined using the Visium Spatial Tissue Optimization Reagents Kit (version 1). Libraries were sequenced on Illumina sequencers (Illumina, San Diego, CA). Cell Ranger (10x Genomics) version 5.0.0 ‘mkfastq’ command was used to generate Fastq files. Space Ranger version 1.2.1 ‘count’ was used with default parameters and alignment to GRCh38 to perform image alignment, tissue detection, barcode and UMI counting, and generation of feature-barcode matrix.

## Author Contributions

N.Z. and H.J. conceived the idea and provided funding support. A.S. performed Visium data generation for patients 4 and 5. C.W. and J.R. performed tumor subclone analysis. P.H. and E.F. performed histopathological analysis for all samples used. K.M developed the method, implemented the software, and performed simulations. K.M. and A.S. analyzed the liver metastasis data. K.M. and N.Z. wrote the manuscript with help from A.S.

